# The hydrocup: a hollow electrospun scaffold for *in vivo* hydrogel-laden cell delivery

**DOI:** 10.64898/2026.09.19.752872

**Authors:** David Boaventura Gomes, Jip Zonderland, Steven Vermeulen, Silvia Rezzola, Ana Henriques Ferreira Lourenço, Denis van Beurden, Jinjie Venema, Hong Liu, Marloes Peters, Peter Emans, Nicole Bouvy, Timo Rademakers, Paul Wieringa, Sandra Camarero Espinosa, Lorenzo Moroni

## Abstract

For regenerative medicine applications, transplanted cells are difficult to retain in a specific place to maximize the localized release of cytokines, which can positively modulate immune responses, angiogenesis and tissue regeneration. Here, we developed a hollow electrospun (ESP) scaffold with one open and one closed end to enclose a hydrogel laden with human mesenchymal stem cells (hMSCs); we termed this combination the hydrocup. hMSCs remained viable for 28 days *in vitro* and released functional cytokines similar to hMSCs in transplanted but non-enclosed hydrogels. The hydrocups remained intact and fixed in place for 6 weeks after subcutaneous implantation in rats, demonstrating *in vivo* compatibility. These results demonstrate that the hydrocup is the first, to our knowledge, cell-laden hydrogel delivery scaffold. This innovation could be developed for a wide variety of cell-based cytokine secretion or drug-laden hydrogel applications.

## Introduction

Cytokines released by human mesenchymal stromal cells (hMSCs) have a wide variety of effects on other cells or local tissues. These effects include immunomodulation, angiogenesis, chemoattraction, anti-scarring and supporting local stem cells in tissue regeneration [1–3]. Indeed, hMSCs are now used in clinical trials as “secretion factories” for their cytokine secretion profile in a wide variety of diseases [4–6]. Searching for “mesenchymal stem cells” or “mesenchymal stromal cells” on clinicaltrials.gov (August 2025) results in 1843 and 428 ongoing or completed clinical trials, respectively, showing the vast amount of ongoing clinical research using MSCs. Many clinical trials use intravenous cell injections. While this route could be useful for systemic release of hMSC cytokines, local implantation can deliver cytokines precisely where they are needed, which could reduce the required cell number and potential adverse effects, such as cells entrapped in capillary beds [7–9]. However, as with any cell transplantation, local cellular delivery and retention in one place remains a challenge [7–12]. For example, direct injection of cells in the target tissue often results in high cell death and low retention [10–12], with one study showing no detectable MSCs 7 days after direct injection into the heart [13]. MSCs and other locally injected cells have been shown to migrate to other tissues throughout the host animal, and disappearing by week 3–4 [14].

We hypothesized that cells could be encapsulated in an electrospun (ESP) scaffold to improve cell survival and retention after implantation. ESP scaffolds are porous and therefore allow for sufficient nutrient or cytokine diffusion, making them good candidates for encapsulated cell delivery [15]. In addition, ESP scaffolds provide a barrier between the surrounding tissue and the implanted cells, which can prevent migration and/or phenotypic changes from contact with other cells. Because MSCs embedded into hydrogels have been shown to exhibit superior survival and retention *in vivo* [13], we embedded the cells first in a hydrogel, which was then enclosed within the ESP scaffold. We developed a cup-shaped ESP scaffold, a hollow electrospun cylinder with one closed end for containment and one open end to enable pipetting in the cell-laden hydrogel (Figure 1). We termed the combination of the cup-shaped ESP scaffold and the hydrogel “the hydrocup.” We experimented with different design parameters, by optimizing the spinning time, including hydrocups with thin (75 µm) and thick (300 µm) walls, to fabricate hydrocups supporting maximal hMSC survival and retention. We demonstrate that hMSCs remained alive in the hydrocups for 28 days and secreted functional cytokines. We then tested the hydrocups for *in vivo* implantation, and found that they remained intact and fixed in place, still containing the hydrogel, after six weeks. These findings demonstrate a novel scaffold for the successful retention of hMSCs that functionally secrete cytokines, which has broad potential for various cell-delivery applications.

**Figure 1.**
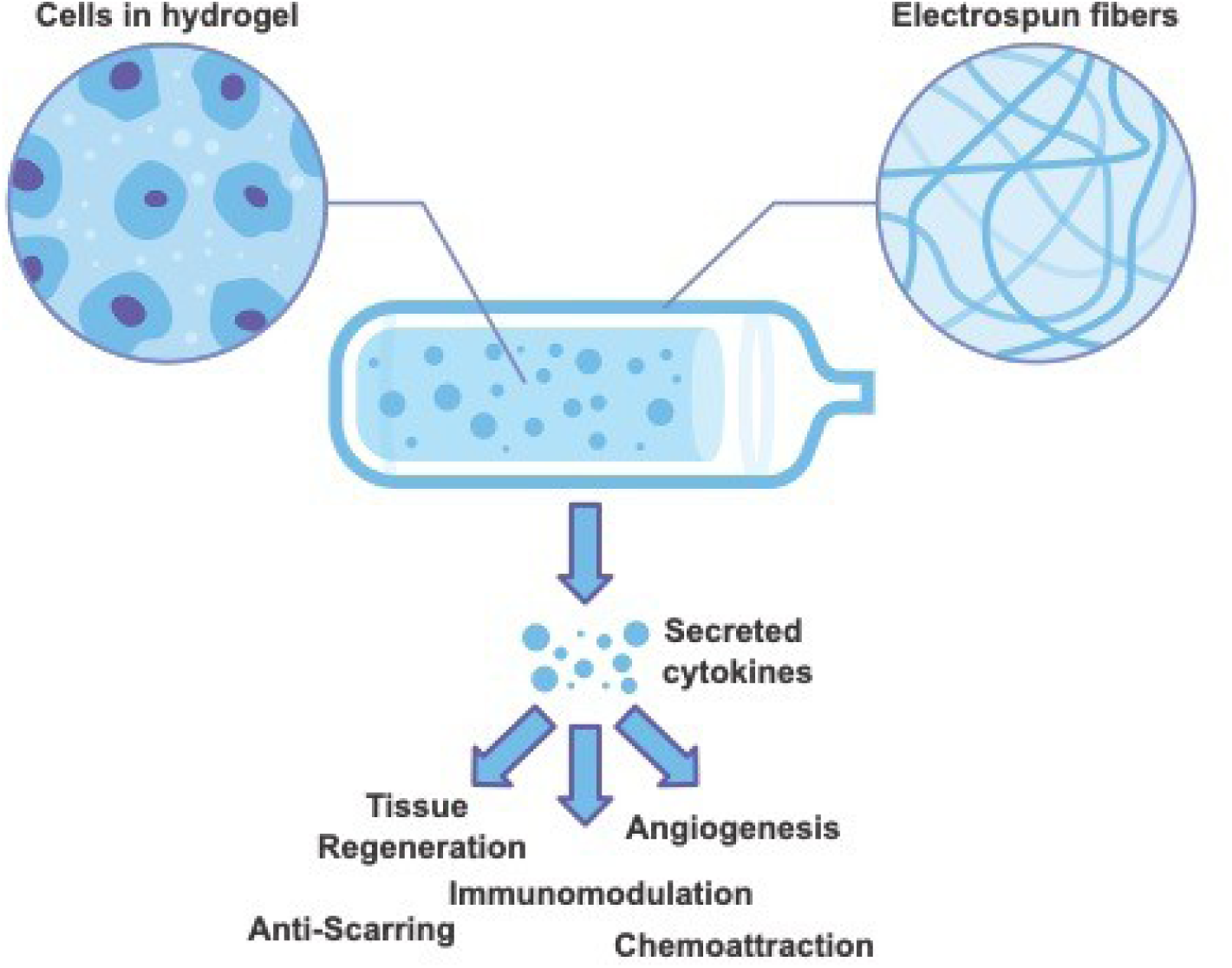
Schematic depiction of the hydrocup, a cell-laden hydrogel encapsulated in an ESP scaffold that could be used to deliver cells *in vivo*.

## Materials and methods

### Hydrocup production

Hydrocups were produced using 300PEOT55PBT45 (PolyVation), synthesized with 300 kDa poly(ethylene glycol), with a 55/45 (w/w) PEOT/PBT ratio. A 35% (w/v) solution of 300PEOT55PBT45 was prepared by overnight dissolution in 30% (v/v) 1,1,1,3,3,3-Hexafluoro-2-propanol AR (HFIP) (Bio-Solve) and 70% (v/v) chloroform (Sigma-Aldrich) under agitation at room temperature (RT). The polymer solution was electrospun on a 1.8 mm diameter mandrel rotating at 100 RPM. The mandrel was attached to the motor on one side and open on the other side. The open side had a rounded tip to prevent charge accumulation at focal points. The polymer solution was fed at 3 ml/h to a charged (10–15 kV) needle in a chamber of 23–25 °C and 30–40% relative humidity. The needle was positioned at an approximately 30° angle to increase coverage on the open end of the mandrel charged to -1.5 kV. After an initial spinning period of ∼1 min in the middle of the mandrel, the needle was moved to the side to spin on the open end of the mandrel. The working distance was approximately 18 cm, depending on the final position of the needle. Thin hydrocups were produced in 30 s and thick hydrocups in 2 min of electrospinning. The hydrocups were removed from the mandrel with 70% (v/v) ethanol, dried and sterilized for 2 h with 254 nm UV light in vacuum. For flat ESP scaffolds, the same conditions were used, but electrospun on a flat collector plate instead of the open-ended mandrel.

### Alginate purification and RGD functionalization

Alginate with 70% “GG blocks” was kindly provided by FMC polymers. Food-grade alginate was purified using a previously published protocol by Neves et al. [16]. Briefly, 1% (w/v) alginate was dissolved overnight under agitation in ultrapure water (18 MΩ, Milli-Q UltraPure Water System, Millipore). To the alginate solution, 2% (w/v) activated charcoal (Sigma-Aldrich) was added and incubated for 1 h under agitation at RT. The suspension was centrifuged at 27 000×*g* for 1 h (Beckman). The supernatant was filtered through a series of different diameter membrane pores (1.2, 0.45 and 0.22 μm, VWR), freeze-dried and stored at −20 °C. To allow cell adhesion, the alginate was modified with (glycine)4-arginine-glycine-aspartic acid-serine-proline (RGD) (Genscript) by using aqueous carbodiimide (EDC) chemistry, as described previously [17]. Briefly, the purified alginate was dissolved at 1% (w/v) in 0.1 M 2-(N-morpholino) ethanesulfonic acid (MES) buffer solution (0.1 M MES buffering salt, 0.3 M NaCl, pH adjusted to 6.5 using 1 M NaOH, Sigma). N-Hydroxy-sulfosuccinimide (sulfo-NHS, Pierce Chemical) and 1-ethyl-(dimethylaminopropyl)-carbodiimide (EDC, Sigma, 27.40 mg per g alginate), at a molar ratio of 1:2, was then added to the solutions, followed by 65.9 μmol RGD for every gram alginate. This mixture was incubated under agitation for 20 h and subsequently quenched with 18 mg of hydroxylamine hydrochloride (Sigma) per gram alginate. The final product was dialyzed over three days at 4 °C (MWCO 3500, Spectra/Por, VWR) against decreasing concentrations of NaCl in ultrapure water, freeze-dried and stored at -20 °C. The RGD-modified alginate is hereafter referred to as “alginate.”

### Cell culture

hMSCs were isolated from the bone-marrow of a 22-year-old male by aspiration, after ethical approval from the local and national authorities and written consent from the donor. The isolation and verification of differentiation capacity was done by Texas A&M Health Science Center [18] and were received at passage 1. hMSCs were further expanded at 1000 cells/cm^2^ in αMEM+Glutamax medium (Thermo Fisher Scientific) supplemented with 10% (V/V) fetal bovine serum (FBS) (Sigma-Aldrich) (basic medium) at 37 °C in 5% CO_2_. Human dermal fibroblasts (Lonza) were expanded at 2000 cells/cm^2^ in DMEM+Glutamax medium (Thermo Fisher Scientific) supplemented with 10% (v/v) FBS (fibroblast medium). At 70–80% confluency, hMSCs and fibroblasts were passaged using 0.05% trypsin and 0.53 mM EDTA (ThermoFisher Scientific) and were used for experiments at passage 5.

### hMSC embedding in alginate

After trypsinization, hMSCs were pelleted by centrifugation at 500×*g* for 5 min and resuspended in 1% alginate (w/v) at 10^6^ cells/ml. For *in vitro* alginate-only conditions (not inserted inside hydrocups), a droplet of 20 µl alginate (containing 2×10^4^ hMSCs) was formed by dispensing the hydrogel with cells on top of a 100 mM CaCl_2_ (Sigma-Aldrich) bath and left to crosslink for 5 min, to be subsequently cultured in basic medium. For *in vivo* alginate-only experiments, 100 µl alginate (containing 10^5^ hMSCs) was crosslinked and implanted. For *in vitro* and *in vivo* experiments with hydrocups, 20 µl alginate (containing 2×10^4^ hMSCs) was dispensed into the bottom of the hydrocup using a 10 µl pipette tip. The hydrocup was directly submerged in a 100 mM CaCl_2_ bath for 5 min. The open end of the hydrocup was then closed with surgical sutures and cultured further in basic medium. Experiments or conditions without cells were performed identically, but without embedded hMSCs.

### Live/dead staining and presto blue assay

To stain live and dead cells inside the hydrocups or alginate-only, they were incubated with 6 µM ethidium homodimer (ThermoFisher Scientific) and 1 µM calcein-AM (ThermoFisher Scientific) for 30 min at 37 °C. Hydrocups or alginate-only were washed 3 times with αMEM without phenol red (Gibco) and imaged directly using an SP8 Leica confocal microscope. Presto blue assay (ThermoFisher Scientific) was performed according to manufacturer’s protocol. Briefly, 14 days after culturing, presto blue reagent was incubated for 2 h at 37 °C with the hMSCs. Fluorescence intensity of the presto blue reagent was subsequently measured at 535/590 nm (excitation/emission) on a ClarioStar plate reader (BMG Labtech, Germany).

### Cytokine assay

To measure cytokine secretion, hMSCs were seeded and cultured for 6 days in 2D tissue culture polystyrene (TCPS) 24-well plates, 3D hydrocups or alginate-only . After this, medium was refreshed with 1 ml basic medium and incubated with the cells for 24 h. In this medium, 105 human cytokines were measured simultaneously using the proteome profiler human XL cytokine array kit (R&D Systems) according to the manufacturer’s protocols. Briefly, the cytokine-containing medium was incubated on PVDF membranes containing spots with antibodies for each cytokine. The provided panel of antibodies was incubated, followed by incubation with horseradish peroxidase–coupled secondary antibodies. Clarity Western ECL (Bio-Rad) was used to visualize the cytokine spots, and all membranes were imaged at the same time to allow relative comparison between conditions.

PCA and hierarchical cluster analysis with complete linkage method were calculated using R version 4.1.2., and visualized with Graphpad Prism version 9.3.1. Prior to both analyses, protein abundance values were mean-centered and scaled to unit variance.

### Migration assay

To test whether functional factors were released by the hMSCs inside the hydrocups to stimulate fibroblast migration, a transwell assay was performed. Fibroblasts (5×10^3^) were seeded on top of a Fluoroblok 24-well transwell with 8 µm pores (Corning) in fibroblast medium. After 6 h of adhesion, the medium in both the top and the bottom compartment was refreshed with fibroblast medium. Alginate-only, thin or thick hydrocups, with or without hMSCs, were added to the bottom compartment. After 24 h, the transwell was washed and fixed with 3.6% (v/v) paraformaldehyde. Fibroblasts were stained with 5 µM Syto14 (ThermoFisher Scientific) and the whole top and bottom of each transwell were imaged. The total number of cells on each side was then quantified under the microscope.

### In vivo subcutaneous implantation

All experiments and protocols were approved by the Dutch Central Committee for Animal Experiments (in Dutch: Centrale Commissie Dierproeven). Female rats aged 8–10 weeks were housed at 21 °C with 12 h light/dark cycles and had *ad libitum* access to water and food (Charles-River, Crl:NIH-Foxn1rnu, 140–212 g). Before anesthesia, 0.05 mg/kg buprenorphine and 4 mg/kg carprofen were administered as premedication. Anesthetization was induced with3–4% (v/v) isoflurane, and maintained with 2% (v/v) isoflurane, and adjusted according to clinical signs during surgery. Four 1 cm long linear skin incisions parallel to the spine were made on the shaved and disinfected animals’ dorsum, two on each side. From these incisions, four subcutaneous pockets (maximum 1×1 cm) were created. Thin or thick hydrocups, or alginate-only, with or without hMSCs, were randomly assigned and placed inside the pockets (n = 8 per condition). The skin was sutured intracutaneously with Monocryl 4x0 sutures (Ethicon) to close the pockets. Approximately 8 hours after surgery, 0.03 mg/kg buprenorphine was administered. On day 1 and 2 post surgery, 4 mg/kg carprofen was given to each animal. Thereafter, animal welfare was evaluated daily. After 6 weeks, CO_2_ overdose was used to euthanize the rats, and sample and surrounding tissues were harvested and processed for histology. No animals were lost during this study.

### Tissue preparation, infiltration quantification and macrophage staining

Tissue explants were fixed for 24 h at 4 °C in a 3.6% (v/v) solution of paraformaldehyde in Tris-buffered saline (TBS) and then underwent embedding in a series of: 30% (w/v) sucrose, then 50:50 (v/v) of 30% sucrose:optimal cutting temperature compound (OCT) (Thermo Fisher Scientific), and then only OCT, for 24 h each. Tissue explants frozen inside silicon molds were filled with OCT on the liquid-vapor interface of a liquid nitrogen tank. Cross-sections of 7 µm were cut on a cryotome onto a film, using the Kawamoto method [19], and samples were stained with safranin (Sigma-Aldrich) O and Fast Green (Sigma-Aldrich) or Masson’s trichrome (Sigma-Aldrich) after hydration in deionized water. For safranin O/Fast Green staining, slides were stained for 3 min in Gill’s haematoxylin (III) (Sigma-Aldrich), washed for 3 min in running water, stained for 3 min with Fast Green, washed in 1% acetic acid for 15 sec, stained 3 min in safranin O. For Masson’s trichrome staining, sections were placed in Gill’s haematoxylin (III) for 5 min, washed with running water for 5 min, dehydrated in ethanol for 15 s and stained with alcoholic Eosin Y (Sigma-Aldrich) for 1 min. For both stainings, sections were then dehydrated in 100% ethanol for 15 s, allowed to air dry and mounted in DPX (Sigma-Aldrich). Assessing macrophage response within the hydrocups was not possible due to the high level of nonspecific binding of the macrophage marker stain 3,3’-diaminobenzidine (DAB) to the hydrocup material itself, resulting in a high level of false-positive cells. Instead, we assessed the macrophage content within a 250 μm wide perimeter around the hydrocups, to investigate how the local environment responds to the hydrocups and their content. Total macrophages (CD68; 1:200, MA513324, ThermoFisher Scientific), M1 (iNOS; 1:300, ab49999, Abcam) and M2 (CD206; 1:100, ab64693, Abcam) were used, diluted in 1% BSA, 4% GS, 0.05% Tween-20 in TBS. Briefly, sections were washed in PBS, after which they were subjected to antigen retrieval: 15 min incubation in 0.05% trypsin/0.1% calcium chloride at 37 °C for CD68 and CD206, and 10 mM citrate buffer, pH 6, for 30 min at 95 °C for iNOS. Thereafter, sections were permeabilized using 0.5% Triton-X100 in PBS, and then blocked using both a peroxidase and protein block (Novolink Polymer Detection System, Leica). Primary antibody staining was performed overnight at 4 °C, after which slides were washed, treated with a post-primary reagent (Novolink Polymer Detection System, Leica) and incubated with the Novolink Polymer for 1 h (Novolink Polymer Detection System, Leica). After washing, slides were developed using DAB for 1 min, and counterstained with haematoxylin. Slides were then mounted by inverting the film containing the section onto a new microscopy slide, using a mixture of 1:1 entallan and xylene. Samples were then dried overnight before imaging. Samples were imaged with an automated inverted Nikon Ti-E microscope, equipped with a Nikon Ri2 sCMOS camera, and an MCL NANO Z200-N TI z-stage, using a 4× objective for imaging of full sections, and 10× or 20× for more detailed images. For imaging full sections, a JOBS protocol was employed to detect the outer boundaries of the tissue section on the slide, after which the system independently mapped the area for stitching and full tissue reconstruction.

### Statistical analysis

Number of replicas is stated in the figure legends. A minimum of n=3 was used for each experiment. Statistical differences were tested using One-way ANOVA with Tukey’s post hoc test. Significance was set at p<0.05. Statistical analysis was performed using GraphPad Prism 9.3.1.

## Results

### Fabricating cup-shaped ESP scaffolds

We fabricated ‘cup-shaped’ scaffolds, with one open end to pipette the alginate in and one closed end to prevent leaking of the uncrosslinked alginate, on the rounded open end of a 1.8 mm rotating mandrel (Figure 2a). By positioning the electrospinning needle at an angle, aiming at the open end of the mandrel, a hollow ESP structure was formed. Using a constant ESP fiber diameter of 2.96 ± 0.24 µm and electrospinning times of 30, 60, 120 and 180 seconds, the wall thicknesses of the resulting hydrocups were ∼75, 150, 300, 450 µm, respectively (Figure 2b). As delamination of the ESP wall occurred in the 450 µm thick hydrocups, subsequent experiments were done with the 75 µm and 300 µm hydrocups, denoted hereafter as ‘thin’ and ‘thick’ hydrocup.

**Figure 2.**
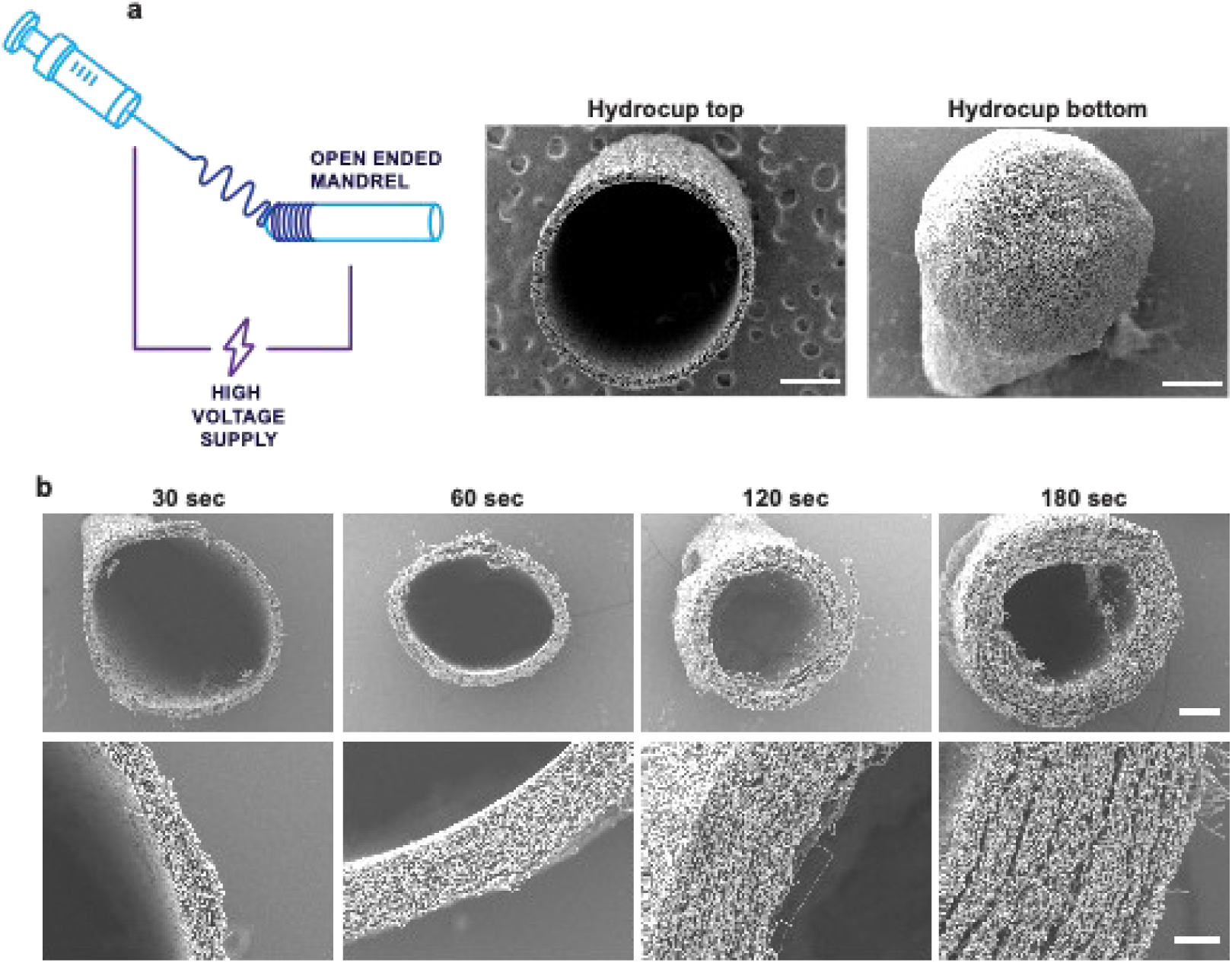
Fabricating ESP scaffolds on an open-ended rotating mandrel. **a**, Schematic depicting the fabrication process of the ESP scaffolds. 300PEOT55PBT45 polymer solution was spun at an angle on the open end of a slowly rotating mandrel for varying durations. After removal from the mandrel, a hollow cylindrical structure with one open end and one closed end was obtained. Scale bars, 500 µm. **b,** Spinning durations resulted in various wall thicknesses of the ESP scaffolds. The top row shows an overview image of the open-ended side of the hydrocup. Scale bar, 500 µm and applies to all panels in the row. The bottom row shows a close-up detail of the ESP wall of the respective image above. Scale bar, 100 µm and applies to all panels in the row.

### Combining hydrogel and ESP scaffold to create the hydrocup

To test whether the alginate could be crosslinked on an ESP scaffold, we crosslinked hMSCs-laden alginate on top of a flat ESP scaffold. The alginate remained in place and attached to the ESP scaffold for 7 days, and hMSCs migrated from the alginate onto the ESP scaffold, together indicating a stable connection between the hydrogel and the ESP scaffold (Supplementary Figure 1). Next, alginate without hMSCs was combined with the cup-shaped ESP scaffolds, hereafter named the hydrocup, to test their initial material binding and compatibility. Alginate was pipetted into the bottom of the ESP scaffold through the open end and directly crosslinked with 100 mM CaCl_2_. The closed bottom of the ESP scaffold prevented leaking of the aqueous, uncrosslinked alginate. Sections were made and alginate was visualized with safranin O staining to investigate if the alginate gel infiltrated the ESP wall of the hydrocups. In both the thin and the thick hydrocups, a thin layer of alginate inside the ESP wall was seen (Figure 3). Alginate did not leak through the hydrocup before crosslinking, preventing loss during the loading process. Also, the layer of alginate in the ESP wall suggests a stable connection between the alginate and the ESP scaffold.

**Figure 3.**
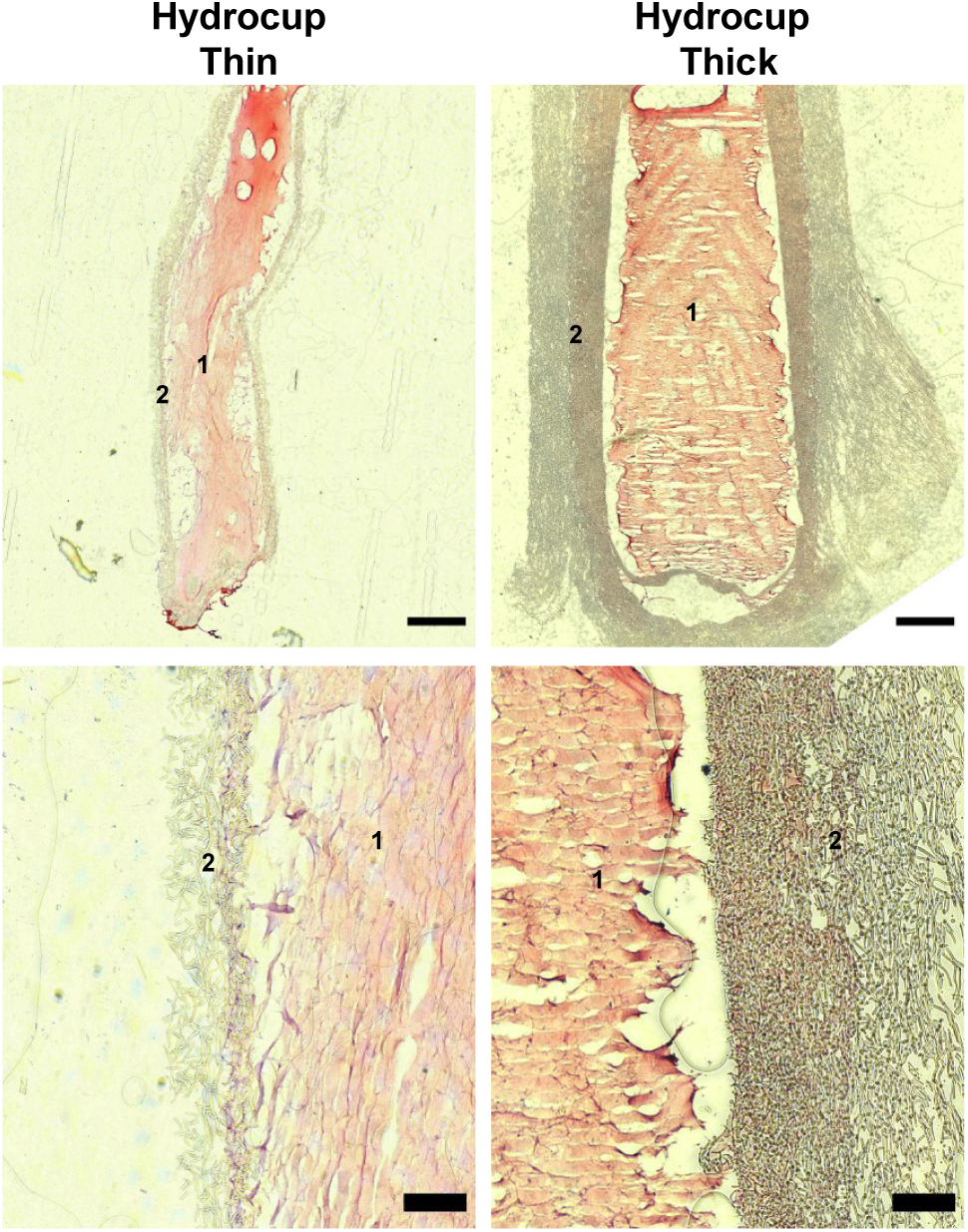
Safranin O staining of thin and thick hydrocups. Sections of hydrocups showing alginate (stained with safranin O, red) (1) (no cells) inside thin (left column) or thick (right column) ESP scaffolds (2). A layer of alginate can be seen in the walls of both the thin and thick hydrocups, suggesting good binding of the alginate and the electrospun wall. The bottom row displays a detail of the wall of the respective image above. Scale bars, 500 µm (top panel) and 100 µm (bottom panel).

### Viable hMSCs contained by hydrocups

To test whether the hydrocups could support cell viability, hMSCs were embedded in alginate that were then loaded into thin and thick hydrocups. The open end of the hydrocups was closed with sutures to contain the cells and alginate. hMSC viability was assessed by live/dead staining at day 7 and day 28 after seeding, and very few dead cells were observed for both time points, with most hMSCs alive (Figure 4a). In addition, a presto blue assay was performed at day 14 to measure cell viability and metabolic activity. No difference was found between hMSCs in thin or thick hydrocups and hMSCs in alginate-only, further confirming that the hydrocups supported high cell viability (Figure 4b).

To determine whether hMSCs migrated outside the hydrocup, the outer walls were imaged at 7 and 28 days post-seeding. On day 7, hMSCs were observed on the outer wall of the thin hydrocup, whereas no cells were observed on the outer wall of the thick hydrocup (Figure 4a). By day 28, hMSCs were present on the outer walls and electrospun fibers of both the thin and thick hydrocups (Figure 4a), which may reflect a combination of cell migration and proliferation. Similar migration was observed for hMSCs encapsulated in alginate on flat ESP scaffolds at day 7 (Supplementary Figure S1). At both day 7 and day 28, live hMSCs were also observed within the interior of the hydrocups. The continued presence of live hMSCs within the hydrogel-filled interior after 28 days indicates that both thin and thick hydrocups are capable of retaining hMSCs in vitro over extended culture periods.

**Figure 4.**
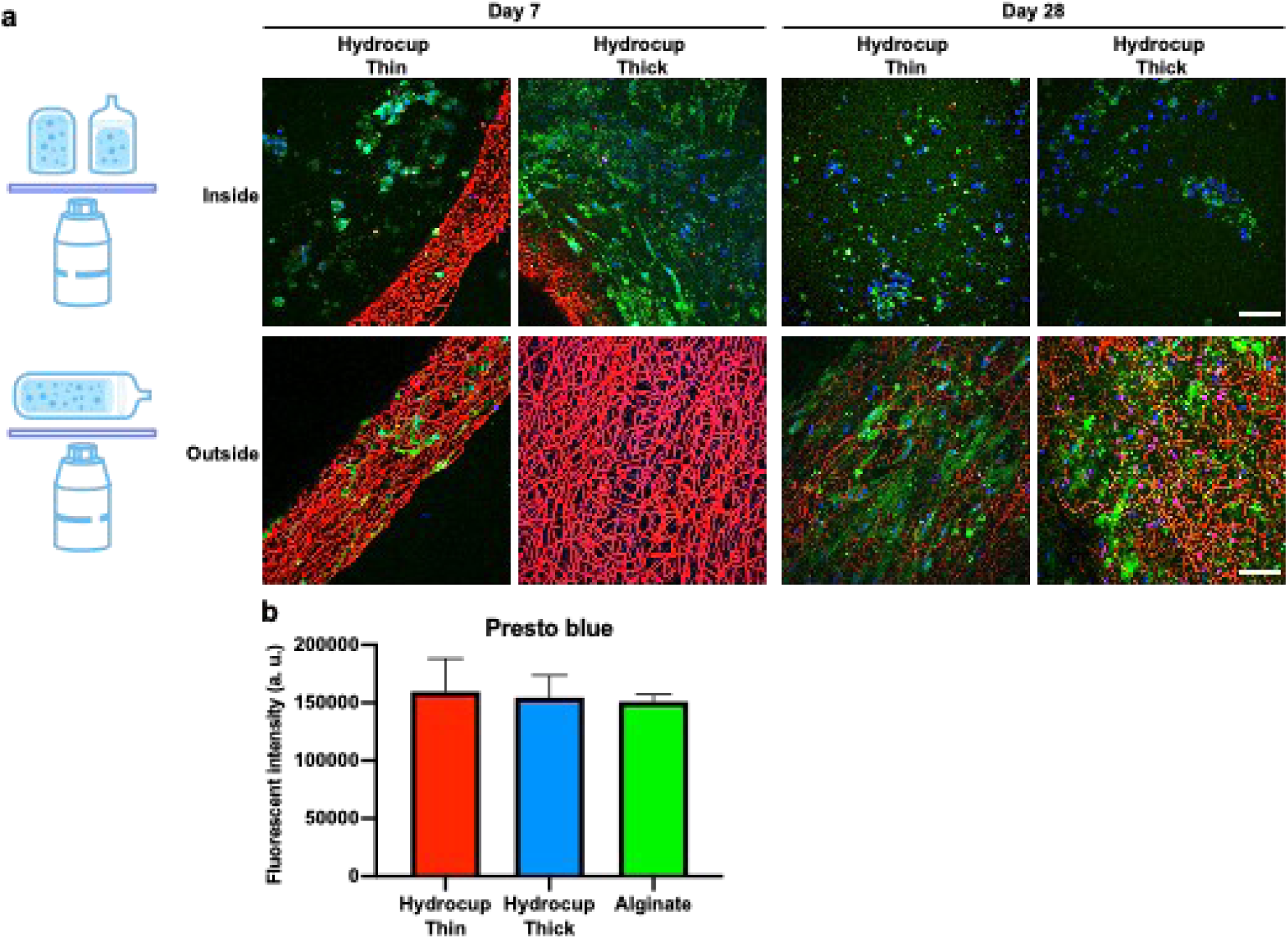
Live/dead staining of hMSCs in thin and thick hydrocups. **a**, Live/dead staining at 7 or 28 days of culture of hMSC-embedded alginate inside the hydrocups. Top panel displays the hMSCs stained inside the hydrocups, while the bottom panel shows hMSCs outside the hydrocups. Green: live cells; blue: nuclei; red: dead cells (inside the hydrocup). The ESP fibers appear as red due to autofluorescence. Representative images of n=3 are shown. Scale bars: 75 µm and applies to all panels of the same row. **b,** Presto blue assay to measure metabolic activity at day 14 after seeding of hMSCs in thin or thick hydrocups, or alginate-only. Bars and error bars are mean ± standard deviations from 3 experiments.

### Functional cytokine release from hMSCs in hydrocup

After determining hMSC viability in the hydrocups, we evaluated their functional activity by profiling the secretion of 105 human cytokines at day 7 post seeding (Supplementary Table S1). As controls, we also tested hMSCs cultured on a flat tissue culture polystyrene (TCPS) and in alginate-alone (not enclosed in the ESP scaffold) at day 7 (Supplementary Figure S2). To evaluate whether the secretion profiles differed between the conditions, we applied principal component analysis (PCA, Figure 5a). We found that the cytokine profile was similar between hMSCs cultured on the thin and thick hydrocup, indicating that the thickness of the ESP scaffold did not meaningfully influence the secretome. However, PCA indicated a distinct cytokine profile between the hydrocups and the alginate-alone, and even more different to TCPS.

In an effort to better understand the unique cytokine profiles elicited by the different culture conditions, we conducted hierarchical cluster analysis, which yielded four major clusters (Figure 5b, Supplementary Figure S3), where the first 3 demonstrated a substantial variety of cytokine secretion across the materials. Of interest, the 4^th^ cluster mainly illustrates that TCPS induced the strongest cytokine secretion into the culture medium, followed by alginate-alone. The average intensity level of all cytokines combined was approximately 5 times higher in the TCPS condition compared to the hydrocups (Figure 5c). A doubling of cytokine intensities was also visible in the alginate-only condition compared to hydrocups. This data indicates that the hMSCs cultured in hydrocups secreted fewer cytokines into the medium compared to alginate-only and classical 2D cell culture conditions.

From clusters 1–3, we identified 3 cytokines that exhibited lower abundance on TCPS compared to alginate and/or the hydrocups (Figure 5d). For chitinase-3-like protein 1 (CHI3L1), all three biomaterials showed elevated levels compared to TCPS (p < 0.001). Similarly, macrophage inhibitory cytokine 1 (MIC1) was elevated on alginate (p < 0.001), hydrocup-thick (p < 0.01) and hydrocup-thin (p < 0.05). For interleukin 8 (IL-8), elevation was observed only on hydrocup-thin (p < 0.05). MIC1 and CHI3L1 were both highest in the alginate alone, and IL-8 was highest from the thin hydrocup (Figure 5d). In cluster 4, the majority of cytokines exhibited elevated abundance on TCPS compared to both alginate and the hydrocups. Of interest, we could identify 4 cytokines that also exhibited significant lower levels in the hydrocups compared to the alginate condition (Figure 4e, Supplementary Tables S2). Interleukin-17A (IL-17A) and neutrophil gelatinase-associated lipocalin (NGAL) were significantly lower in both hydrocup conditions compared to alginate (p < 0.001 for IL17, p<0.05 for NGAL). For osteopontin (OPN), a significant difference compared to alginate was observed only for the thick hydrocup (*p* < 0.01). Similarly, for endothelial plasminogen activator inhibitor (SERPIN E1), a significant difference was observed only for then hydrocup (*p* < 0.05).

Next, we determined whether the secreted cytokines were functional. Fibroblasts were cultured on top of a transwell with an hMSC-laden hydrocup or alginate-alone in the bottom compartment. Indeed, cytokines released from the hMSCs contained by either the thin or the thick hydrocup attracted fibroblasts to the bottom side of the transwell, to a similar extent as hMSCs in alginate-only (Figure 5f). This experiment demonstrates that the secretomes of hMSCs effectively elicited a response from other cells, regardless of the ESP scaffold wall.

**Figure 5.**
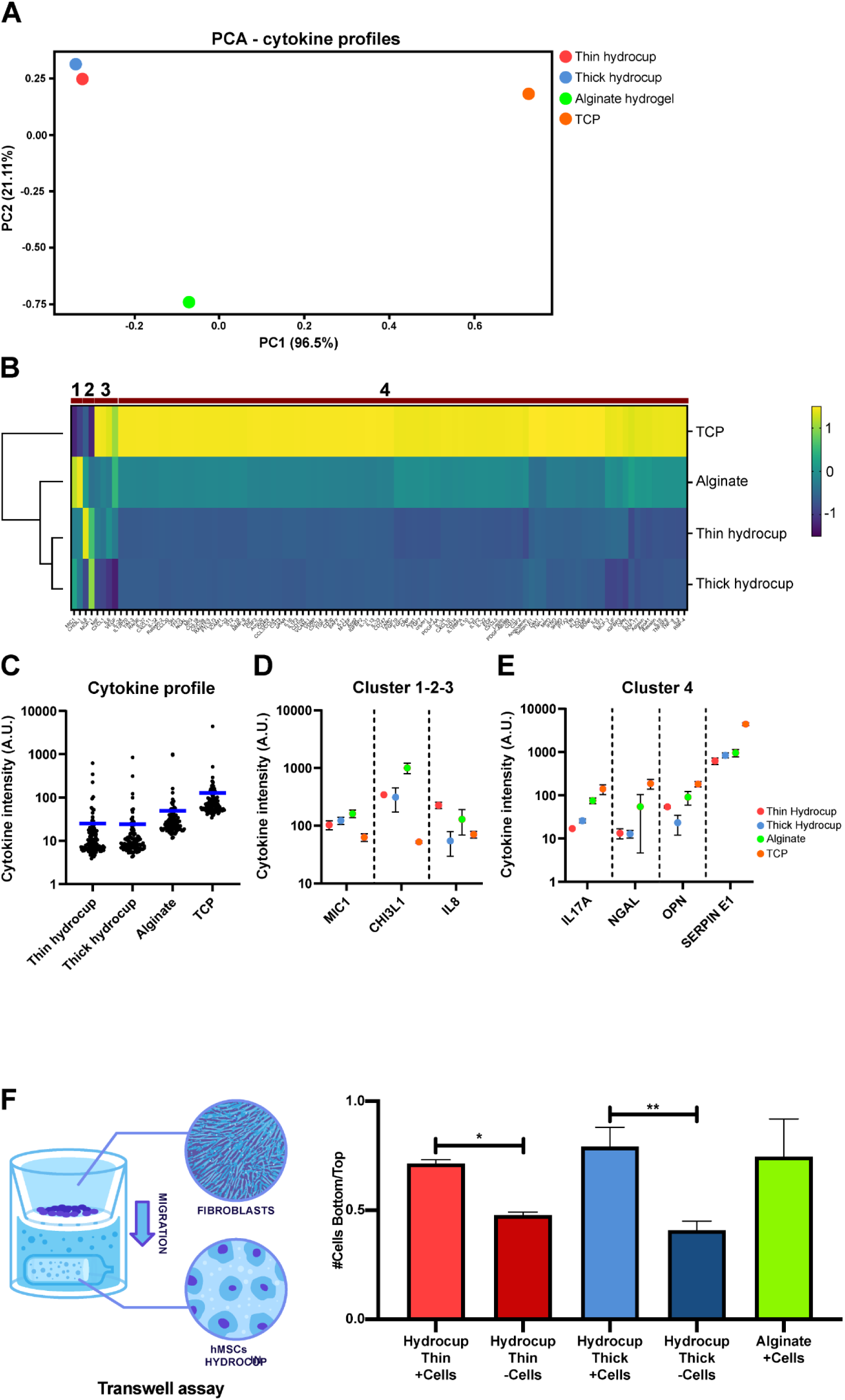
Cytokine secretion of hMSCs inside the hydrocups compared to controls. a–e, hMSCs were cultured for 7 days inside thin or thick hydrocups, alginate alone (without scaffold), and TCPS. Cytokine secretion was measured over the last 24 h, n=3. **a,** PCA analysis of the cytokine secretion profiles demonstrates that the thin and thick hydrocups have a similar cytokine profile, and distinct from the alginate-alone and TCPS conditions. **b,** Hierarchical cluster visualization reveals three small clusters of cytokines with distinct secretion levels across the conditions (cluster 1-2-3), while cluster 4 reveals that the majority of cytokines are elevated in TCPS compared to the hydrocups. **c,** Dotplot visualization of cytokine intensity levels across the conditions shows 5× elevated expression with TCPS culture compared hydrocups. Blue lines represent the average of each measured cytokine. **d,** CHI3L1 was significantly elevated on alginate, hydrocup-thick and hydrocup-thin compared to TCPS (*p* < 0.001). MIC1 was significantly elevated on alginate (*p* < 0.001), hydrocup-thick (*p* < 0.01) and hydrocup-thin (*p* < 0.05). IL-8 was significantly elevated only on hydrocup-thin compared to TCPS (*p* < 0.05). **e**, IL-17A was significantly lower on both hydrocup-thick and hydrocup-thin compared to alginate (both *p* < 0.001). NGAL was significantly lower on both hydrocups compared to alginate (*p* < 0.05). OPN differed significantly between hydrocup-thick and alginate (*p* < 0.01). SERPIN E1 differed significantly between hydrocup-thin and alginate (*p* < 0.05). **f,** Quantification of migrated fibroblasts (ratio from the top and bottom of transwells) after 24 h with alginate-only, thin or thick hydrocups (with or without hMSCs embedded after 7 days) in the bottom compartment. One-way ANOVA; * p<0.05, ** p<0.01. Error bars indicate mean ± SD.

### *In vivo* implantation of hydrocups

To test if the thin and thick hydrocups (with and without embedded hMSCs) could be used *in vivo*, they were implanted in subcutaneous pockets in rats. Alginate-only, also with and without hMSCs, was tested for comparison. The hydrocups were found intact 6 weeks after implantation, demonstrating that the hydrocups can be used to fix hydrogels in place *in vivo*. Both the thin and the thick hydrocups still contained the enclosed alginate, with and without cells.

Sections were prepared and stained for safranin O to stain the alginate and counter-stained with Fast Green to visualize the surrounding tissue. As expected, the alginate was in full contact with the surrounding tissue, sometimes with tissue infiltration throughout the alginate (Figure 6). In the conditions with hMSCs, cells were visible inside the alginate-only and the hydrocup conditions; no cells were found inside the alginate in the conditions without cells. This difference suggests that the observed cells inside the alginate were the originally embedded hMSCs. In the thin hydrocups (with and without cells), *de novo* tissue formed, as seen by Fast Green staining, in the ESP hydrocup wall (Figure 6b, d). A thin layer of *de novo* tissue was also observed in the ESP wall of the thick hydrocups, but the majority of the cup remained free of tissue. This finding suggests that these cells are mostly rat cells infiltrating the hydrocup. Alginate was in direct contact with rat tissue with alginate-only and the thin hydrocup, whereas the alginate in the thick hydrocups remained more isolated (Figure 6b, d). Collagen and tissue formation in the hydrocups was confirmed with Masson’s trichrome staining (Supplementary Figure 4). Collagen capsule formation was observed in all conditions, including alginate, hydrocup thin, and hydrocup thick groups, both with and without cells. These findings indicate that the presence of hMSCs did not interfere with the host response or with collagen capsule formation around the biomaterials.

**Figure 6.**
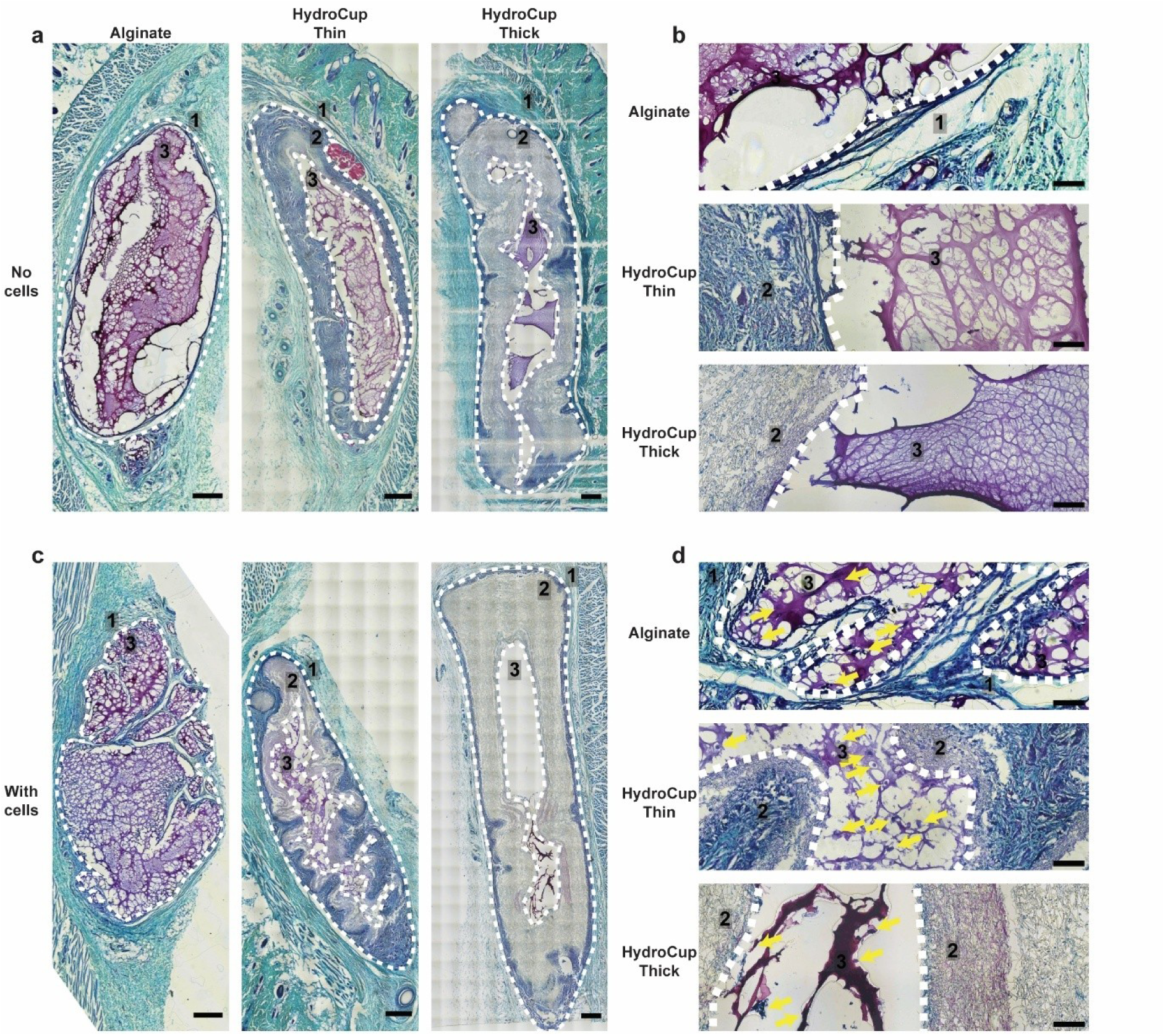
*In vivo* implantation of thin and thick hydrocups compared to alginate-only, with (c, d) or without (a, b) hMSCs embedded. Safranin O/Fast Green staining of sections from 6-week post-subcutaneous implantation in rats. Numbers label the surrounding rat tissue (1), hydrocup ESP wall (2), alginate gel or lumen of the hydrocup (3). **b, d,** detailed images showing changes around the hydrocup or alginate-only. Representative images of n=8 per condition are shown. Scale bars in **a, c**: 500 µm; in **b, d**: 100 µm. Yellow arrows represent cells encapsulated in the hydrogel. Dotted lines delineate the limits of the implanted material.

Finally, we evaluated the inflammatory response to *in vivo* hydrocup implantation, by quantifying macrophages, M1-and M2-type responses, using markers C68, iNOS and CD206, respectively (Figure 7). We observed no significant differences between the thin and thick hydrocups in the number of macrophages in a 250 μm wide perimeter surrounding the hydrocup (Figure 7a) or in M1 and M2 phenotypes (Figure 7b, c). Together, these results indicate that the hydrocups could be fixed in place *in vivo* and remained intact, retaining the cell-laden hydrogel for 6 weeks, and without eliciting substantial immune responses. In addition, the thick hydrocups showed limited tissue formation and protected the embedded alginate from direct contact with surrounding rat tissue.

**Figure 7.**
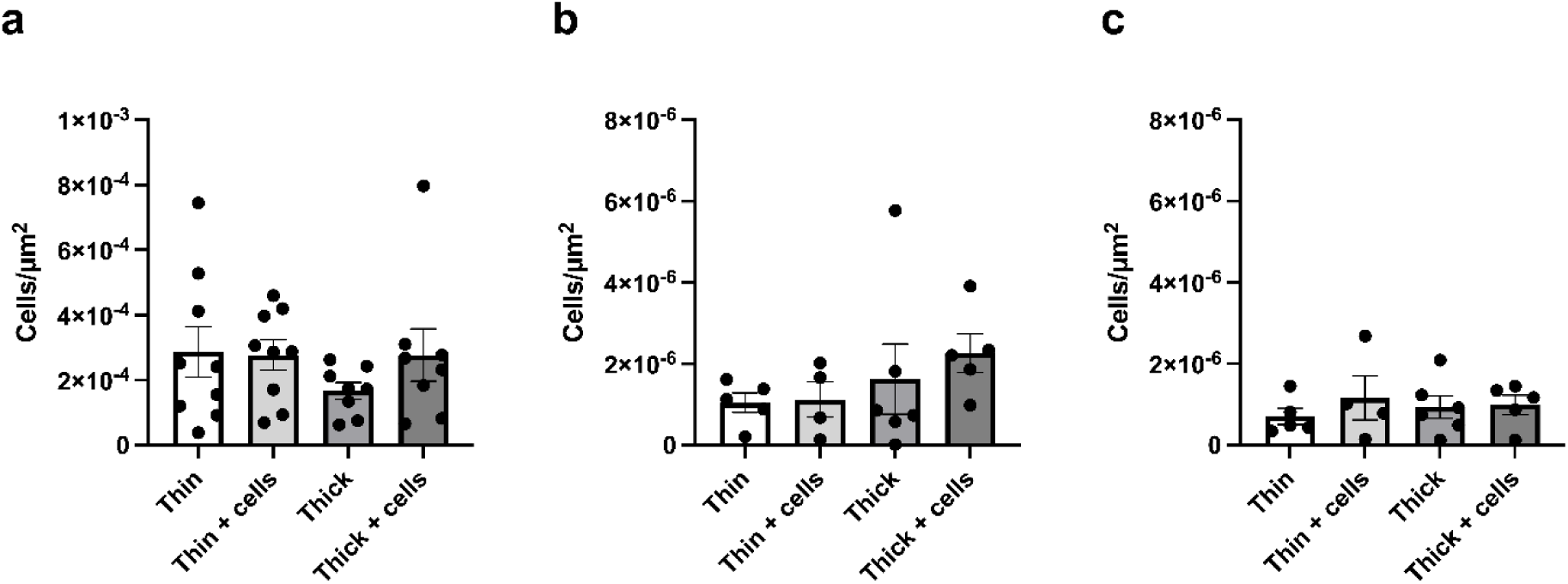
Immune responses from *in vivo* implantation of thin and thick hydrocups, with and without hMSCs embedded. Staining for CD68 (a) showed no significant difference in total macrophage numbers. Additionally, no differences could be observed for M1 (iNOS, b) and M2 (CD206, c) types, respectively. Bars and error bars are mean ± SD from at least 4 experiments.

## Discussion

The hydrocup described here, featuring a tubular ESP scaffold with one closed end and one open end has, to our knowledge, not yet been reported. The formation of tubes with two open ends by electrospinning on a small mandrel has been done for many years, mainly for nerve conduits or vascular grafts [20, 21]. The closed end represents a technological challenge due to the difficulty of controlling fiber deposition in a specific location, on the open end of a thin mandrel. This closed end facilitates hydrogel loading into the scaffold, preventing the watery, uncrosslinked alginate from directly leaking out of the scaffold. The elegance of our approach is that the properties of the hydrocup can easily be adjusted. For example, other polymers could be used to tweak the mechanical or degradation properties, or to promote different interactions with the hydrogel or surrounding tissue. Also, we have made the hydrocups on a 1.8 mm diameter mandrel, but other dimensions could be selected. Additionally, the ≈3 µm fiber diameter used here provided sufficient porosity in the scaffolds for hMSCs to migrate. Smaller fiber diameters could be used to decrease porosity of the ESP scaffolds and limit hMSC migration. Fiber diameters of ≤1 µm could also decrease the infiltration of surrounding tissue cells into the hydrocup wall. While we used hMSCs and alginate here as a proof-of-concept, other cell types or hydrogel materials could be used. Besides cell-laden hydrogels, drug-releasing hydrogels could also be fixed in place using hydrocups.

hMSCs enclosed in the hydrocups remained viable for 28 days (Figure 4) and showed intact signaling as evidenced by cytokine expression (Figure 5) and fibroblast migration (Figure 6). Interestingly, we found distinct cytokine expression profiles between our 3D hydrocups, 3D alginate-only and 2D TCPS conditions, with 2D TCPS being the most different from the other conditions (Figure 5a). Furthermore, the expression level of cytokines detected from hMSCs cultured in 2D TCPS was five times greater than the other conditions, while cells in alginate-only also showed two times more secretion than those in the hydrocups (Figure 5c). These findings align with previously reported observations for hMSCs cultured in 3D scaffolds. The reduced cytokine secretion observed in 3D conditions is consistent with the well-documented phenomenon of cellular quiescence induced by alginate-based matrices with low RGD content, which promotes a metabolically less active state and consequently diminishes both transcriptional activity and protein synthesis [22]. In contrast, 2D monolayer cultures, such as TCPS, support greater cell proliferation and metabolic activity, resulting in higher RNA and protein output. These differences in paracrine signaling between 2D and 3D culture systems have been extensively reported in the literature, with multiple studies demonstrating that the dimensionality of the culture environment profoundly influences the secretory behavior of hMSCs [23, 24]. These differences may be the result of sequestering hMSCs in the closed hydrocups, using these materials as barriers. Such a barrier could also entrap cytokines within the alginate/hydrocup, which would not be captured by our cytokine assay that measured secretion into the medium.

*In vivo* implantation of the hydrocups demonstrated successful retention of the hydrocup and integrity of the enclosed alginate, up to the 6 weeks of observation (Figure 6). Both alginate-only and the hydrocup resulted in the host tissue forming a collagenous capsule. Future studies could tune the material outer layer to one that induces less capsule formation and more vessel formation to a better tissue integration, without compromising the Hydrocup integrity.

While scaffolds have been developed to hold cell aggregates for *in vivo* purposes [25–28] and ESP scaffolds for *in vitro* uses [29], the hydrocup is the first reported scaffold for *in vivo* hydrogel delivery. Delivering cells in a hydrogel rather than directly attached to an ESP scaffold can be beneficial, because the concentration of cells can be far greater within a hydrogel. Also, MSCs in alginate hydrogels have shown superior survival and retention after *in vivo* implantation than directly injected MSCs [13]. The hydrocups could be used to further increase this retention and survival, allowing for the hydrogel to be easily sutured in place and potentially aid cell-based “secretory factory” approaches. Hydrocups can be used in cases such as heart tissue, muscle, or joints, where hydrogels are difficult to place due to movement. In such an approach, cytokine release could potentially be steered by changing the hydrogel properties; the alginate used here was functionalized with the RGD motif but many other possibilities are available. Future studies could explore the vast customization properties and *in vivo* potential of the novel hydrocups reported here.

## Conclusions

We have developed the hydrocup: a hollow cylindrical ESP scaffold with an open-and a closed end for the *in vivo* delivery of hydrogels. hMSCs inside the hydrocups maintained high viability for at least 28 days *in vitro* and the cytokine release profile of hMSCs in the hydrocups was largely similar to hMSCs in alginate only. The hydrogel-containing hydrocups remained intact and fixed in position for 6 weeks *in vivo*.

## Acknowledgements

We thank David Koper for his help with the surgeries and Guilherme Gomes for helping create the illustrations in the figures. This research has been made possible by the European Research Council starting grant “Cell Hybridge” for financial support under the Horizon2020 framework program (Grant #637308), Dutch Research Council (NWO, Grant #16711) and the Dutch Province of Limburg (LINK project). Some materials used in this work were provided by the Texas A&M Health Science Center College of Medicine Institute for Regenerative Medicine at Scott & White through a grant from NCRR of the NIH (Grant #P40RR017447).

## Conflicts of Interest

The authors declare no conflict of interests.

## Supplementary Information

## Supplementary Information

**Supplementary Table S1.** List of cytokines, chemokines, and acute phase proteins detected in a single sample by the Proteome Profiler Human XL Cytokine Array Kit (R&D systems).

|  |  |  |
| --- | --- | --- |
| Adiponectin/Acrp30 | IFN-gamma | CCL2/MCP-1 |
| Angiogenin | IGFBP-2 | CCL7/MCP-3 |
| Angiopoietin-1 | IGFBP-3 | M-CSF |
| Angiopoietin-2 | IL-1 alpha/IL-1F1 | MIF |
| Apolipoprotein A1 | IL-1 beta/IL-1F2 | CXCL9/MIG |
| BAFF/BlyS/TNFSF13B | IL-1ra/IL-1F3 | CCL3/CCL4 MIP-1 alpha/beta |
| BDNF | IL-2 | CCL20/MIP-3 alpha |
| CD14 | IL-3 | CCL19/MIP-3 beta |
| CD30 | IL-4 | MMP-9 |
| CD31/PECAM-1 | IL-5 | Myeloperoxidase |
| CD40 Ligand/TNFSF5 | IL-6 | Osteopontin (OPN) |
| Chitinase 3-like | IL-8 | PDGF-AA |
| Complement Component C5/C5a | IL-10 | PDGF-AB/BB |
| Complement Factor D | IL-11 | Pentraxin 3/TSF-14 |
| C-Reactive Protein/CRP | IL-12 p70 | CXCL4/PF4 |
| Cripto-1 | IL-13 | RAGE |
| Cystatin C | IL-15 | CCL5/RANTES |
| Dkk-1 | IL-16 | RBP4 |
| DPPIV/CD26 | IL-17A | Relaxin-2 |
| EGF | IL-18 BPa | Resistin |
| CXCL5/ENA-78 | IL-19 | CXCL12/SDF-1 alpha |
| Endoglin/CD105 | IL-22 | Serpin E1/PAI-1 |
| EMMPRIN | IL-23 | SHBG |
| Fas Ligand | IL-24 | ST2/IL1 R4 |
| FGF basic | IL-27 | CCL17/TARC |
| KGF/FGF-7 | IL-31 | TFF3 |
| FGF-19 | IL-32 alpha/beta/gamma | TfR |
| Flt-3 Ligand | IL-33 | TGF-alpha |
| G-CSF | IL-34 | Thrombospondin-1 |
| GDF-15 | CXCL10/IP-10 | TIM-1 |
| GM-CSF | CXCL11/I-TAC | TNF-alpha |
| CXCL1/GRO alpha | Kallikrein 3/PSA | uPAR |
| Growth Hormone (GH) | Leptin | VCAM-1 |
| HGF | LIF | VEGF |
| ICAM-1/CD54 | Lipocalin-2/NGAL | Vitamin D BP |

**Supplementary Table S2.**
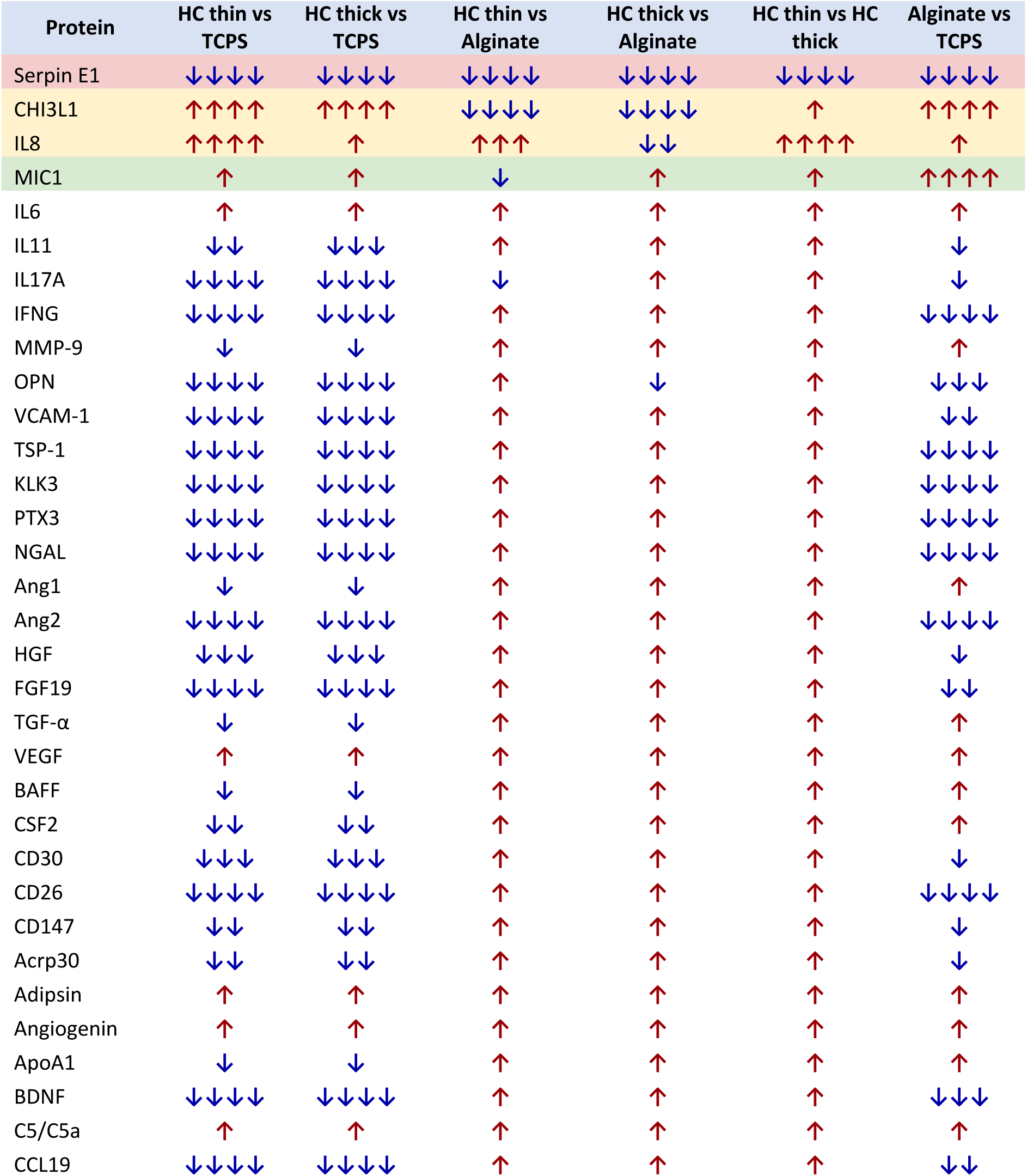
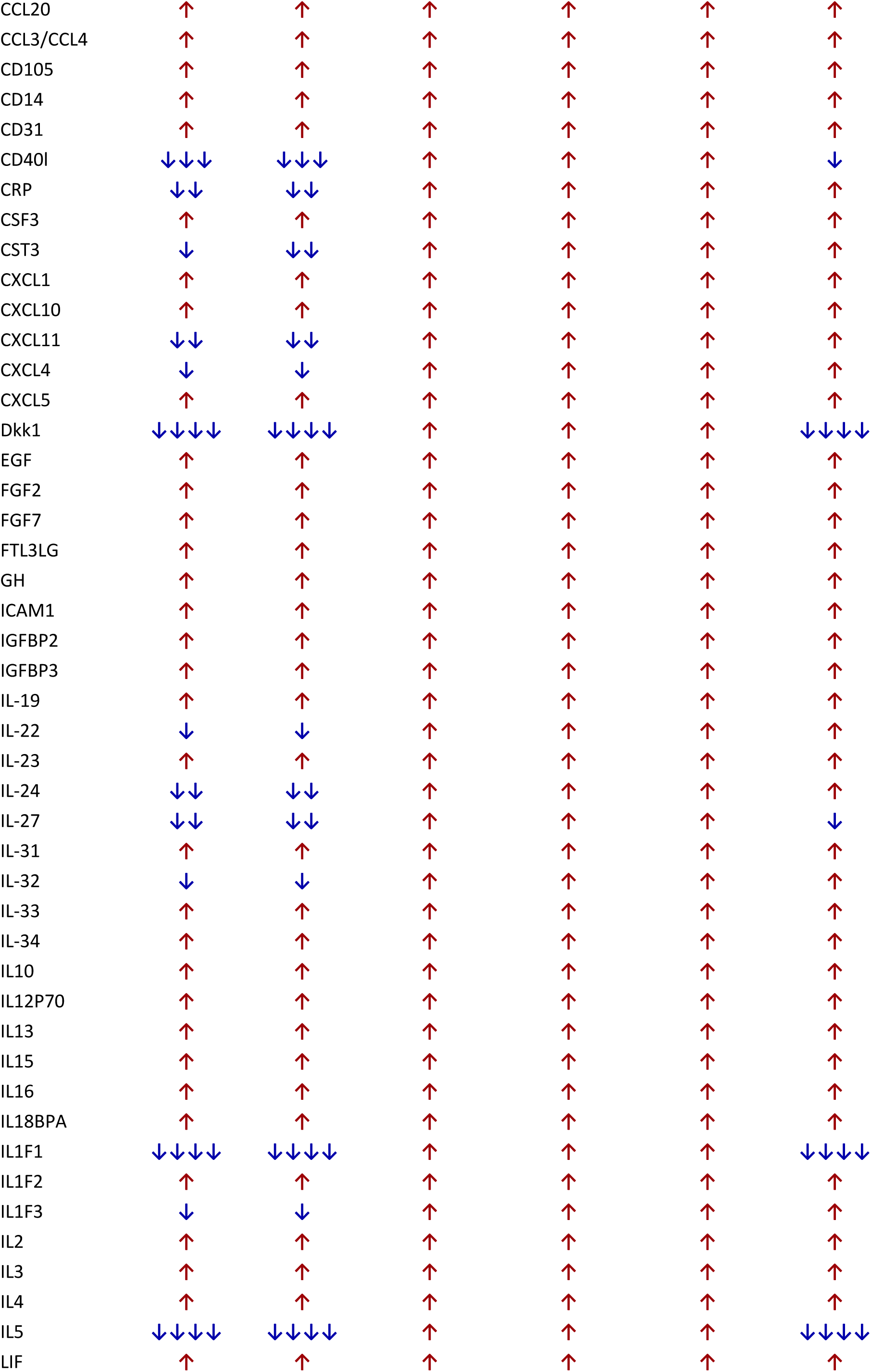
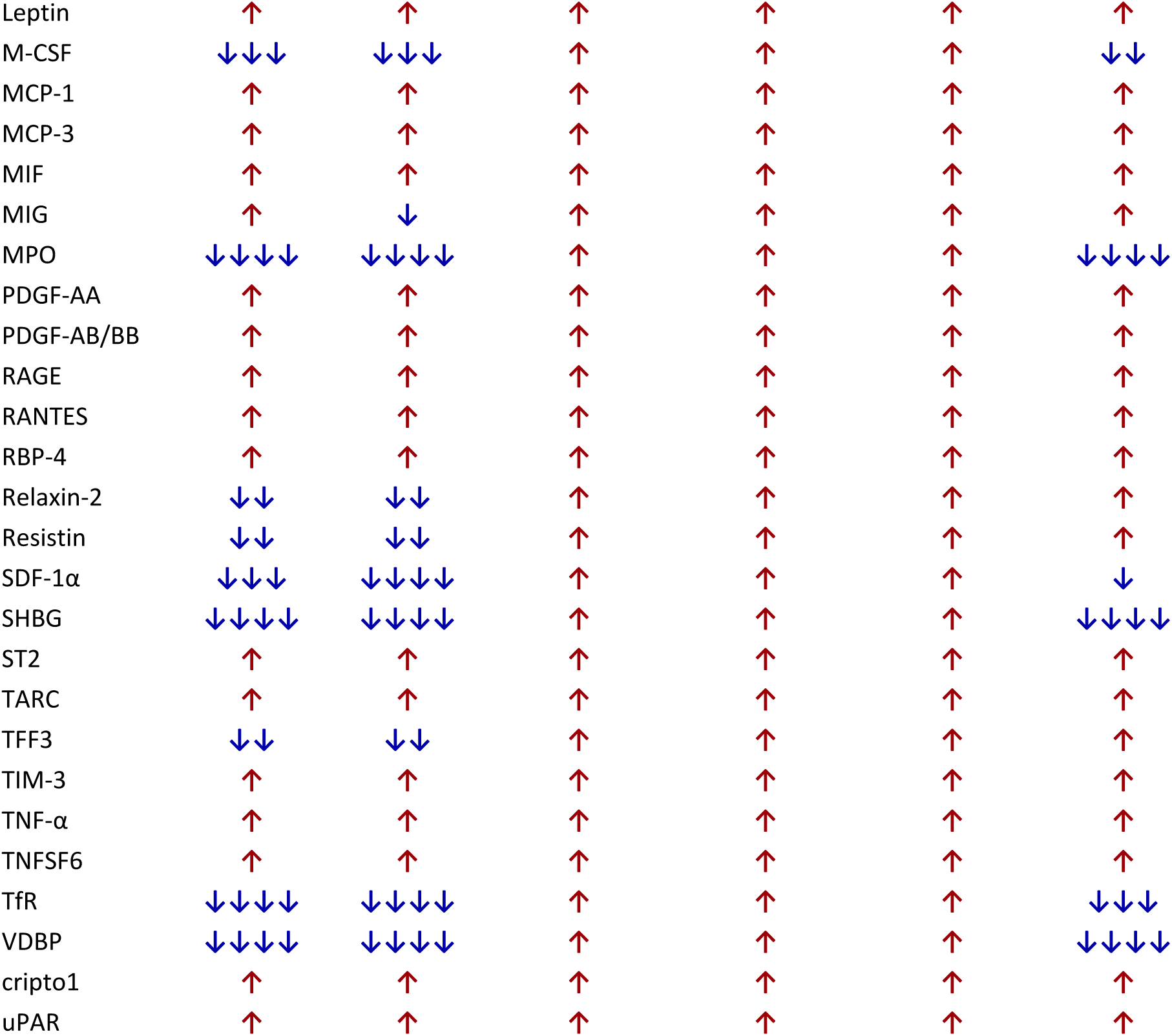
Differential secretome profiles across TCPS, alginate, HydroCup thin, and HydroCup thick conditions. Pairwise comparisons of significantly altered proteins from the 105-analyte secretome panel are shown. Arrows indicate increased (↑) or decreased (↓) secretion relative to the comparator group, with arrow number corresponding to statistical significance (p < 0.05, *p < 0.01, **p < 0.001, ***p < 0.0001). Most significant differences were observed between TCPS and hydrogel conditions, whereas HydroCup thin and HydroCup thick showed largely similar secretion profiles. Serpin E1 was significant across all pairwise comparisons, while CHI3L1 and IL-8 most strongly differentiated hydrogel conditions. Statistical analysis was performed using one-way ANOVA with Tukey’s multiple-comparisons test.

**Supplementary Figure S1.**
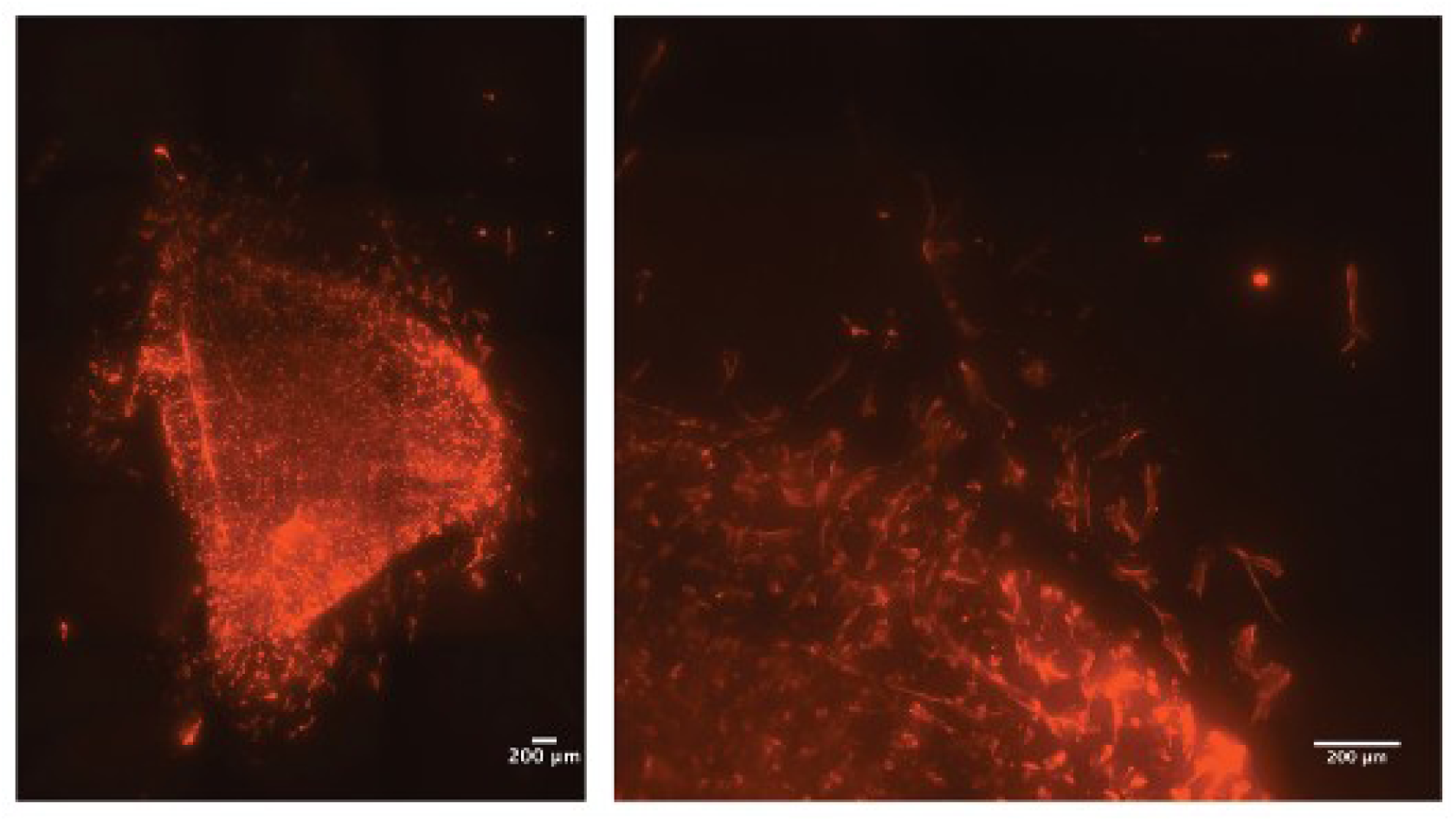
Combination of alginate and flat ESP scaffold. hMSCs embedded in alginate were crosslinked on top of a flat ESP scaffold. At day 7, hMSCs migrated from the hydrogel to the ESP scaffold in the overview (left) and zoom (right). Scale bars, 200 µm.

**Supplementary Figure S2.**
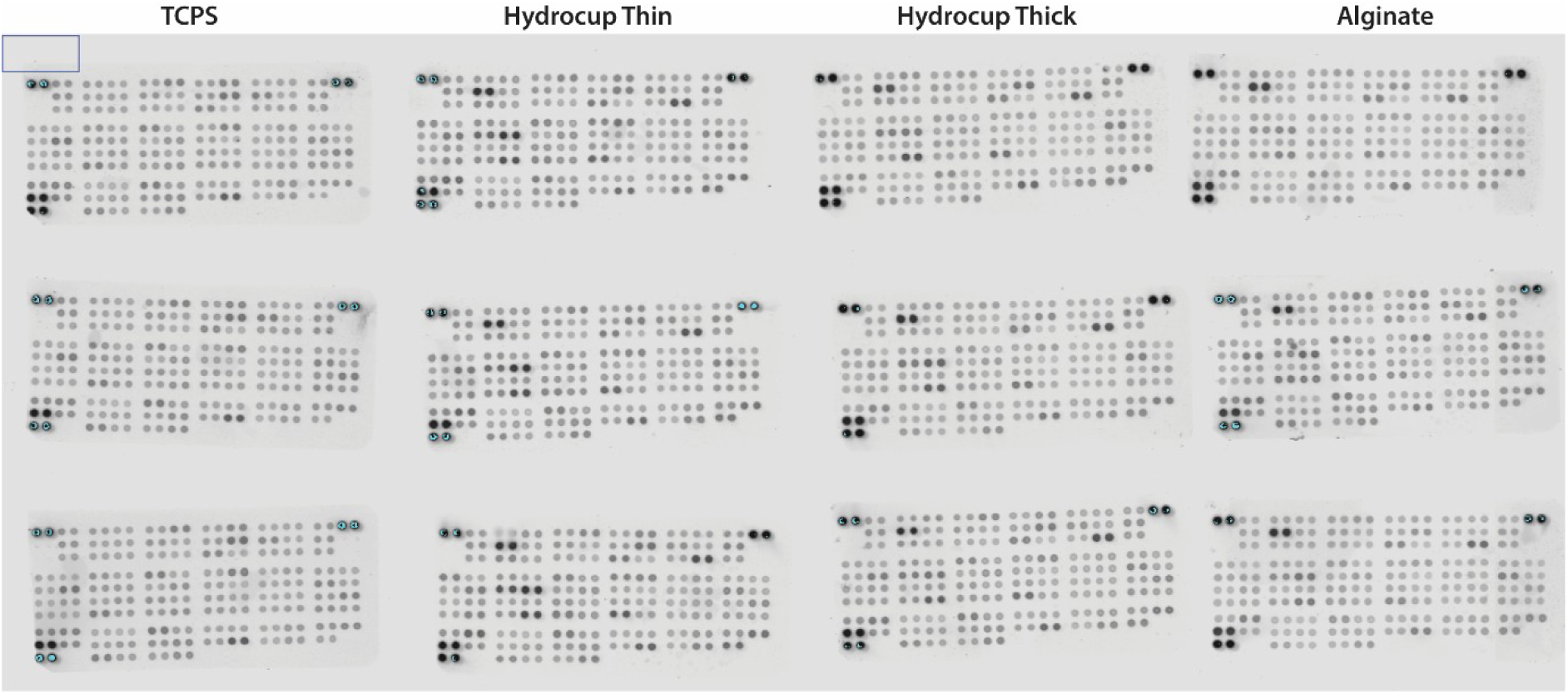
Blots of cytokine secretion in the culture medium taken at 7 days postseeding from hMSCs in thin or thick hydrocups or in alginate-only. Blots were spotted with antibodies against 105 different cytokines. Each condition was done in triplicate and imaged at the same time to allow for relative comparison between conditions.

**Supplementary Figure S3.**
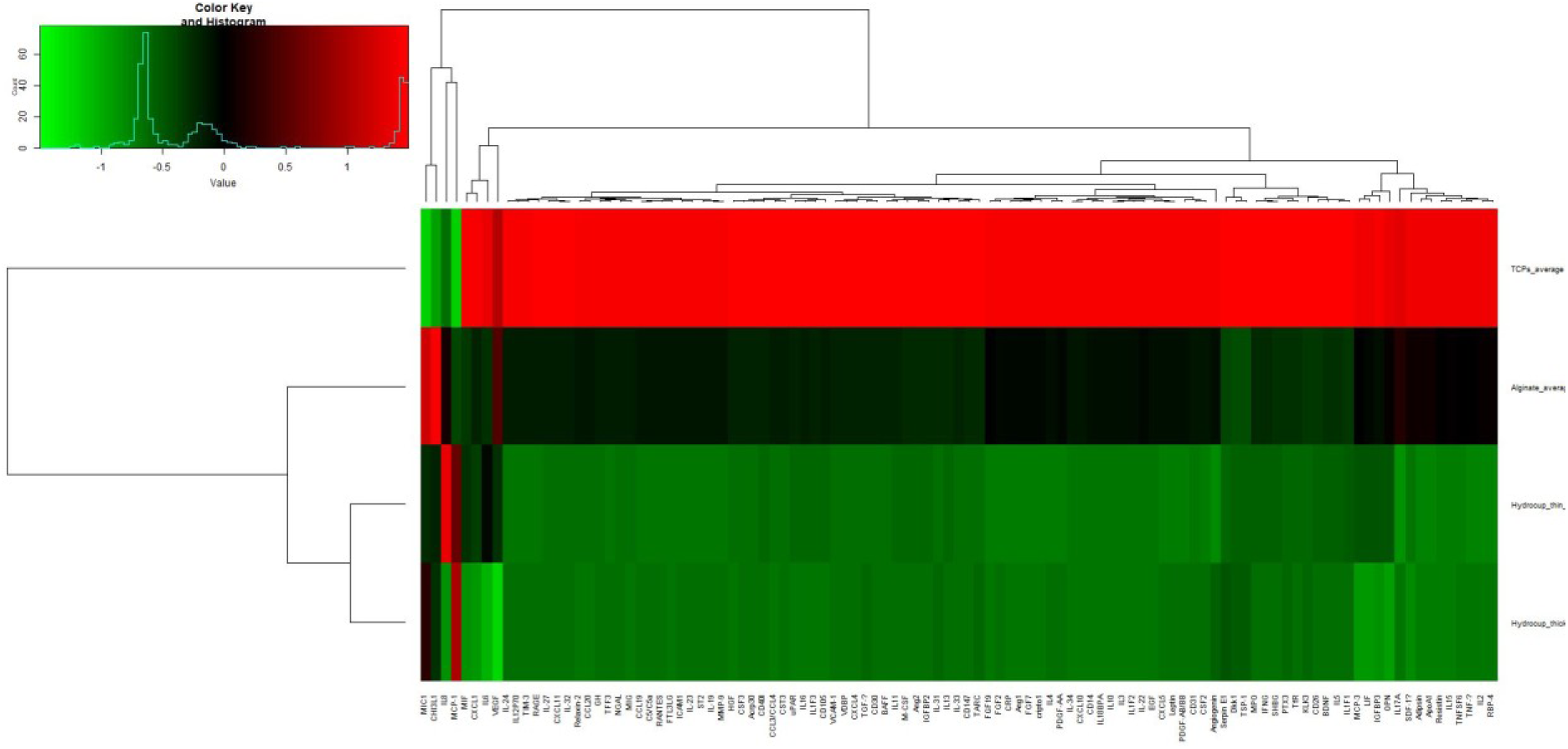
Original hierarchical clustering heatmap of the secreted proteome of hMSCs cultured on 2D tissue culture polystyrene (TCPS), 3D alginate-alone, 3D thin and thick hydrocups. Each column represents an individual secreted protein and each row represents the culture conditions, averaged across replicates. Colour encodes the normalized secreted level of each protein. Red indicates higher secreted levels and green indicates lower secreted levels.

**Supplementary Figure S4.**
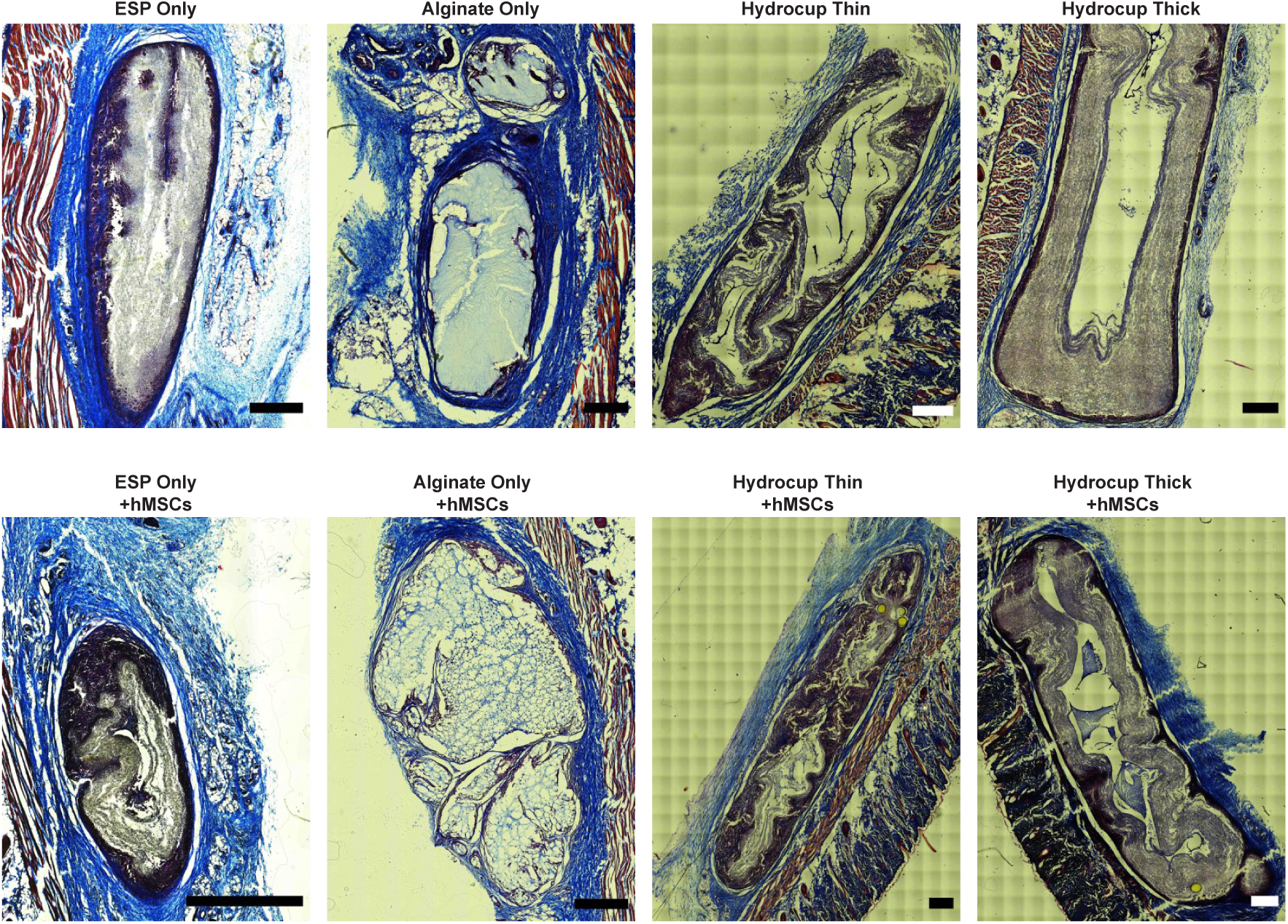
Masson’s Trichrome staining of *in vivo* implanted thin and thick hydrocups, alginate-only and ESP only, without (top row) or with hMSCs (bottom row). Scale bars, 500 µm.

## Notes

### Competing Interest Statement

The authors have declared no competing interest.

